# Low levels of H5N1 HA and NA antibodies in the human population are boosted by seasonal A/H1N1 infection but not by A/H3N2 infection or influenza vaccination

**DOI:** 10.1101/2025.07.13.664638

**Authors:** Anne P. Werner, Cosette G. Schneider, Elgin H. Akin, Juliahna Hayes, Katherine Z. J. Fenstermacher, Richard E. Rothman, Lynda Coughlan, Andrew Pekosz

**Affiliations:** W. Harry Feinstone Department of Molecular Microbiology and Immunology, The Johns Hopkins Bloomberg School of Public Health, 615 North Wolfe Street, rm W2116, Baltimore, MD, 21205, USA; Department of Microbiology and Immunology, University of Maryland School of Medicine, Baltimore, MD 21201, USA; Department of Emergency Medicine, Johns Hopkins University School of Medicine, Baltimore, MD, USA; Center for Vaccine Development and Global Health (CVD), University of Maryland School of Medicine, Baltimore, MD 21201, USA

## Abstract

An increase in the number of human cases of influenza A/H5N1 infection in the US has raised concerns about the pandemic potential of the virus. Preexisting population immunity is a key determinant for risk assessment and pandemic potential for any virus. Antibody responses against the bovine A/H5N1 hemagglutinin (HA) and neuraminidase (NA) proteins were measured among a population of influenza-vaccinated or influenza-infected individuals. Modest titers of bovine A/H5N1 HA-binding antibodies and low to undetectable neutralizing antibody responses were detected in a cohort of 73 individuals. Conversely, bovine A/H5N1 NA binding and neuraminidase-inhibiting antibody responses were comparable to those against a human A/H1N1 NA at baseline. Seasonal influenza vaccination failed to significantly increase antibody titers against both HA and NA glycoproteins of bovine A/H5N1. Recent infection with human A/H1N1 but not A/H3N2 viruses induced significant increases in bovine A/H5N1 neutralizing antibody, as well as increases in NA-binding and NA-inhibiting antibodies to bovine A/H5N1 NA. While the degree of protection afforded by these A/H5N1 cross-reactive antibodies is not known, incorporating NA or enhancing current seasonal vaccine formulations to increase NA-specific antibody responses may increase antibody breadth and protection against both seasonal and pandemic influenza viruses.

## INTRODUCTION

Since its arrival in North America in late 2021, highly pathogenic avian influenza (HPAI) A/H5N1 clade 2.3.4.4b has been a major threat to human and animal health^1–3^. Despite only sporadic outbreaks in humans, with nearly all cases directly traceable to exposures to infected animals, and the lack of evidence of human-to-human transmission, the high case fatality rate (CFR) of roughly 52% seen globally for related H5N1 clades substantiates significant concern over its pandemic potential^1,3,4^.

The A/H5N1 virus has spread to a large number of migratory birds and that has led to hundreds of spillovers into a range of wild mammals, avian species such as birds of prey (not usually hosts for avian influenza viruses), domesticated poultry, household cats, and dairy cows. The A/H5N1 clade 2.3.4.4b viruses have also undergone reassortment with other avian influenza viruses, leading to a number of different genotypes including B3.13 (responsible for most dairy cow infections) and D1.1 (driving a large number of poultry farm infections^5^.

As of June, 2025, the US Centers for Disease Control report 70 cases of A/H5N1 infection, primarily in individuals working in dairy or poultry farms with known exposures to infected animals^2^. While one death has been reported, the vast majority of cases have resulted in relatively mild infection, often limited to conjunctivitis^6–8^. This is in contrast to both outbreaks of A/H5N1 HPAI in prior years and contemporary outbreaks outside of North America ^9–11^. For example, of the 10 reported cases of A/H5N1 in Cambodia in 2025 alone, 6 of them have been fatal^11,12^. Sporadic spillovers have occurred to humans with no evidence of human-to-human transmission^1,3,4^. A/H5N1 viruses isolated from a subset of these infections have shown amino acid changes consistent with adaption to replication in mammals, which highlights concerns about the virus evolving into a form that may lead to a pandemic^13–15^.

Because A/H5NX (where X is one of a number of neuraminidase subtypes) influenza viruses have not circulated in the human population, the vast majority of humans lack antibody responses against the H5 hemagglutinin (HA) protein—the major antigenic target of the influenza immune response. However, human seasonal A/H1N1 viruses encode a neuraminidase (NA) protein that is antigenically and structurally similar to that of A/H5N1 viruses. Experiments in animals and surveys of human cohorts have indicated that antibodies to seasonal human A/H1N1 NA can cross-react with the A/H5N1 NA, but it is unknown if these low-level cross-reactive responses can provide protection against or mediate severity of A/H5N1 HPAI infection in humans^16–19^.

To determine the level of preexisting human population immunity to bovine-derived A/H5N1 viruses, serum samples from two distinct cohorts – healthcare workers vaccinated with the 2024 seasonal influenza vaccine and individuals infected with seasonal A/H1N1 or A/H3N2 influenza during the 2023-24 influenza season – were tested for the presence of antibodies recognizing the H5 HA or N1 NA proteins. Functional antibody responses—neutralizing antibody (nAb) and neuraminidase-inhibiting (NAI) antibody— against bovine A/H5N1 HA and NA were quantified in both cohorts. Low levels of binding IgG to the bovine H5 HA correlate with minimal-to-undetectable nAb responses to bovine A/H5N1 viruses, and responses are not boosted by standard seasonal vaccination. Similarly, binding IgG and NAI antibody responses to human seasonal A/H1N1 NA and bovine A/H5N1 NA were quantified and were comparable at baseline (i.e., prior to vaccination or infection). Increases in NA-specific antibody to both human and bovine N1 NAs following infection with seasonal A/H1N1 viruses, but not A/H3N2 viruses were detected. Lastly, despite no detectable change in N1-binding responses post-A/H3N2 infection, baseline serum depletion of A/H3N2 NA-specific antibody reduces total bovine N1 binding IgG, suggesting that heterotypic NA antibody may play a role in baseline cross-reactivity. Our findings corroborate and extend existing evidence that current seasonal vaccine formulations are poor at inducing NA-specific humoral responses^18,20^.

## RESULTS

### Study design and demographics

Population-level antibody responses to A/H5N1 HPAI neuraminidase (NA) in healthy adults was investigated utilizing a cohort of individuals employed by the Johns Hopkins Hospital (JHH) or Johns Hopkins Medical Institutes (JHMI) who were receiving their annual dose of trivalent 2024-2025 Northern Hemisphere (NH) seasonal inactivated influenza vaccine (IIV) in September and October 2024. Vaccinees provided blood samples at the time of enrollment, indicated as “day 0,” or “baseline,” and at day 28 post-vaccination (**Supplemental Figure 1a**). 50 participants were selected, comprised of 25 males and 25 females, with ages ranging between 23 years to 68 years (**Table 1**). 47 of 50 (94%) participants received egg-grown vaccine (Fluarix, GSK), and 3 participants (6%) whom had reported egg allergy and were immunized with standard trivalent cell-grown vaccine (Flucelvax, Seqirus) (**Table 1**). Only 2 of 50 (4%) participants reported that they had not received any influenza vaccination in any of the past five NH influenza seasons (**Table 1**). Of the remaining 48 (96%) participants, the average number of previous vaccinated seasons was 4.46/5.

**Table 1.**
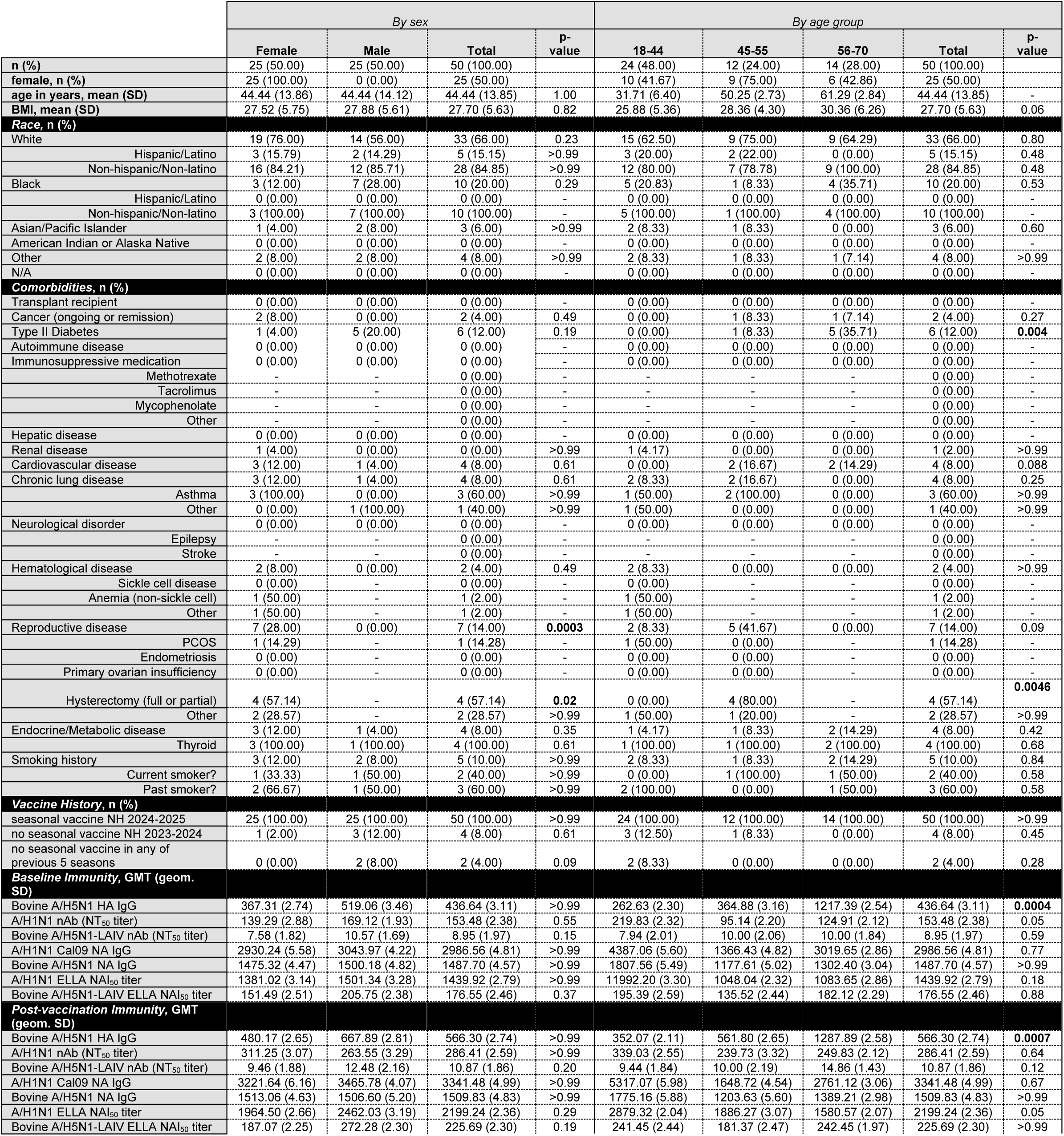
Vaccine Cohort characteristics, demographics, and geometric mean titers for all assessments described. For qualitative data and co-morbidities, Fisher’s exact test was used to generate p-values shown. For baseline and post-vaccination immunity readouts, adjusted p-values shown were calculated via Kruskal-Wallis nonparametric test with Bonferroni’s correction for age analyses, and Dunn’s test for multiple comparisons for sex analyses, both with Bonferroni’s corrections. Bolded p-values are significant (i.e., p < 0.05). Percentages are calculated as total of parent category.

To investigate the role of seasonal IAV infection in shaping cross-reactive antibody responses to bovine A/H5N1 HA and NA, a cohort of patients who presented to the JHH Emergency Department (JHH-ED) or were inpatients at the JHH with influenza-like illness (ILI) with confirmed seasonal IAV infection during 2023-24 Northern Hemisphere influenza season were utilized. IAV infection was confirmed via point of care diagnostic tests or next-generation sequencing. All 23 patients provided blood samples at the time of admittance, hereafter referred to as “baseline,” and again at approximately 4 weeks later, hereafter referred to as “convalescent” (**Supplemental Figure 1b**). 16 of 23 (69.6%) patients were infected with A/H1N1pdm09-like viruses, and 7 (30.4%) with A/H3N2-like viruses (**Table 2**). 14 of 23 (60.9%) participants were female and 9 of 23 (39.1%) were male (**Table 2**). 3 patients (13%) were solid organ transplant recipients, and on immunosuppressive medication (**Table 2**). 2 patients (8.7%) had previously undergone chemotherapeutic cancer treatment and were in remission at the time of the study (**Table 2**). 13 of 23 (56.7%) patients had reported receipt of a seasonal influenza vaccine for the 2023-2024 NH season (**Table 2**). For co-morbidities, 5 patients (21.7%) reported cardiovascular disease, 5 patients (21.7%) reported hematological disorders, 2 patients (8.7%) reported renal disease, and 3 patients (13%) reported asthma (**Table 2**).

**Table 2.**
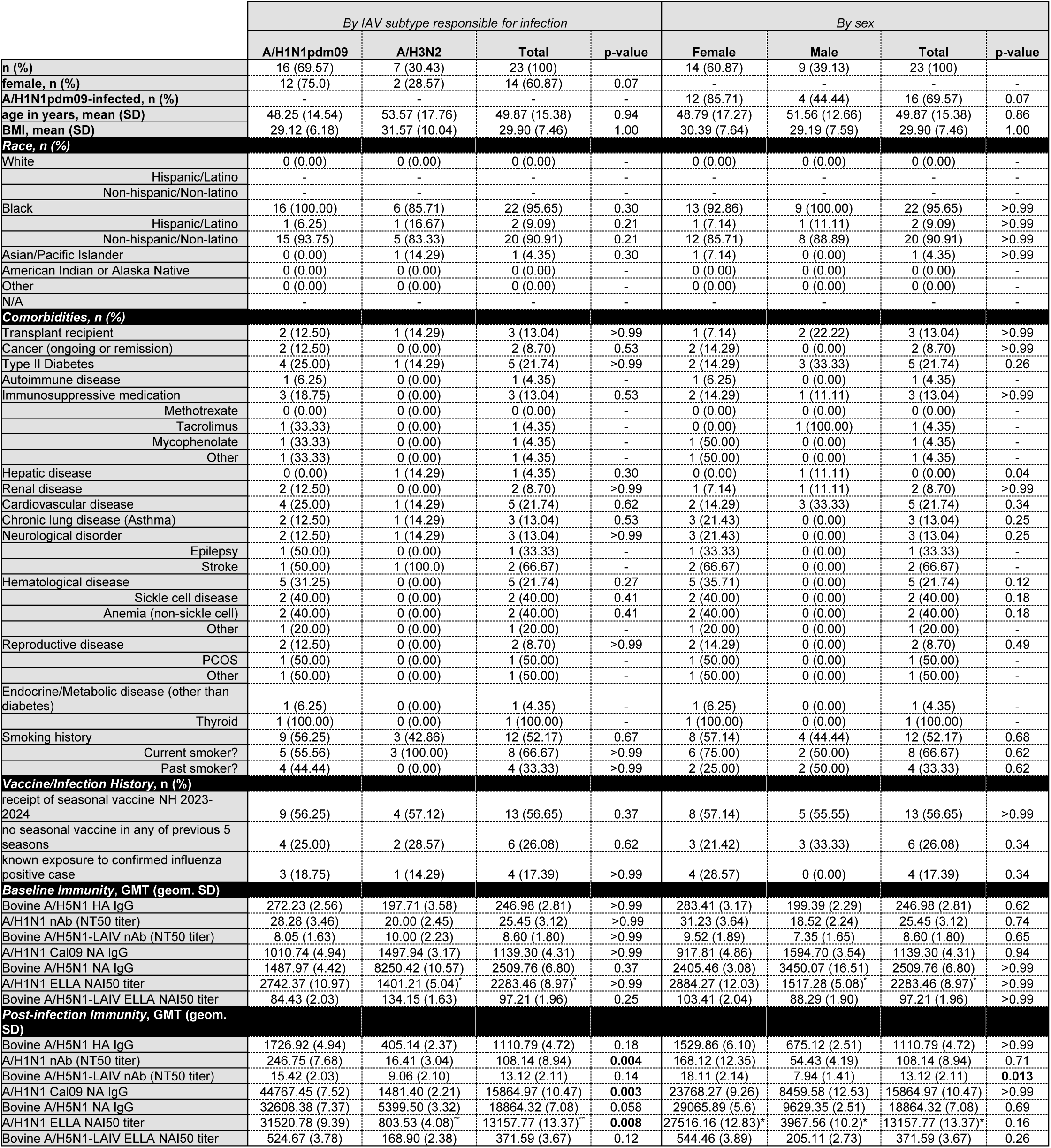
Infection Cohort characteristics, demographics, and geometric mean titers for all assessments described. For qualitative data and co-morbidities, Fisher’s exact test was used to generate p-values shown. For baseline and post-vaccination immunity readouts, adjusted p-values shown were calculated via Dunn’s test for multiple comparisons with Bonferroni’s correction. Bolded p-values are significant (i.e., p < 0.05). Percentages are calculated as total of parent category. Percentages are calculated as total of parent category. *one sample of the indicated group was not run due to limited sample volume, **two samples of the indicated group were not run due to limited sample volume.

### Baseline cross-reactive antibody responses to bovine A/H5N1 NA

To capture the range of baseline cross-reactivity across both cohorts, serum samples at the time of enrollment for the vaccine cohort and at the time of patient testing for the infection cohort were used to determine total binding IgG, NA inhibiting (NAI) antibody, and neutralizing antibody (nAb) responses (**Figure 1**). Baseline IgG binding responses are highly comparable against the bovine N1 NA and the NA from A/California/04/2009 (Cal09), which represents a human seasonal A/H1N1 pandemic09-lineage NA. All participants had detectable binding IgG against the bovine and Cal09 NA at baseline, with GMTs of 1,754.17 and 2204.47 respectively (**Figure 1a**). Although anti-Cal09 NA-binding titers were significantly correlated with birth year, this trend was less clear for anti-bovine N1 NA IgG (**Figure 1a**). Baseline binding titers against both Cal09 NA and bovine A/H5N1 NA were significantly higher than baseline binding titers to bovine H5 HA GMT = 364.88 (**Supplemental Figure 2a**). This suggests that pre-existing antibody responses to bovine N1 NA are greater than to bovine H5 HA.

**Figure 1.**
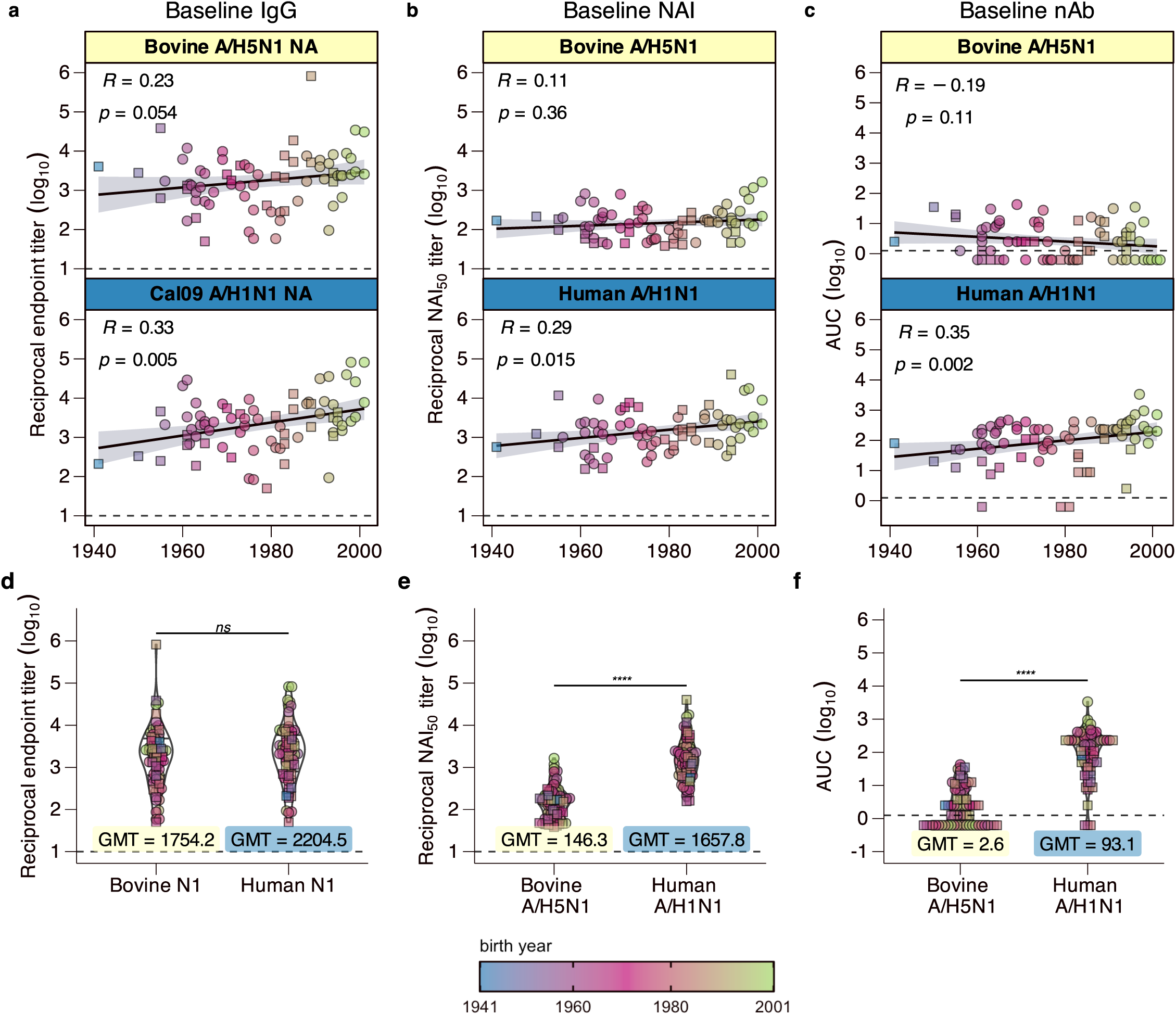
Baseline cross-reactive responses against A/H5N1. (n = 73 total at baseline timepoint; squares represent participants in the infection cohort and circles represent participants in vaccine cohort) Baseline blood serum was used to investigate baseline responses against A/H1N1 NA and A/H5N1 NA. (**a, d**) ELISA binding IgG against the bovine A/H5N1 NA (top panel, yellow label), and against a representative human seasonal A/H1N1 NA, from A/California/04/2009 (bottom panel, blue label), plotted by (**a**) birth year and (**d**) antigen. (**b, e**) Enzyme-linked lectin assay (ELLA) NA inhibition (NAI) responses at baseline against an A/H5N1 bovine recombinant virus in an LAIV backbone (top panel, yellow label) and against a contemporary A/H1N1 seasonal virus (bottom panel, blue label), plotted by (**b**) birth year and (**e**) virus. (**c, f**) neutralizing antibody (nAb) titers at baseline against an A/H5N1 bovine recombinant virus in an LAIV backbone (top panel, yellow label) and against a contemporary A/H1N1 seasonal virus (bottom panel, blue label), plotted by (**c**) birth year and (**f**) virus. (**a-c**) spearman correlation coefficient and significance are indicated in the upper lefthand corner of each panel. (**d-f**) box-and-violin plots of the data shown in a-c with geometric mean titers (GMTs) indicated for each (**d**) antigen or (**e-f**) virus. (**a-f**) Dotted lines indicate the lower limit of detection for the specified assay. (**a-c**) Line of best fit is indicated by the black line, with the standard error indicated by the shaded area in each panel. (**d-f**) Significance shown were generated via nonparametric one-way ANOVA with Dunn’s post-test. *ns = not significant*, *\*\*\*\* = p < 0.0001*.

Using a recombinant bovine A/H5N1 virus whose multibasic cleavage site was deleted (ΔMBS) and was expressed in the genetic background of a live-attenuated influenza vaccine virus (LAIV), hereafter referred to as bovine A/H5N1-LAIV, NAI responses were evaluated using enzyme-linked lectin assay (ELLA). Unlike binding IgG, baseline NA-inhibiting (NAI) responses against the bovine N1 NA were markedly lower than those against a human seasonal N1 NA (**Figure 1b**).

Bovine A/H5N1-LAIV nAb responses were significantly lower than nAb responses against human A/H1N1 (**Figure 1c**), with 46% of subjects (24 of 50 vaccinees and 10 of 23 infected participants) having no detectable neutralizing antibody responses at baseline (**Figure 1c**). We detected a strong correlation between baseline H5-binding IgG and both participant birth year (**Supplemental Figure 2a**) and bovine A/H5N1 nAb response (**Supplemental Figure 2c**). Conversely, only 0.04% of participants (3 of the 23 infected participants and none of the 50 vaccinees) had no neutralizing responses to A/H1N1 at baseline (**Figure 1c**). Taken altogether, pre-existing immunity at the population level to bovine A/H5N1 HPAI preferentially targets N1 NA, rather than H5 HA.

### Seasonal vaccination does not boost cross-reactive bovine A/H5N1 responses

Seasonal vaccines are currently the only widely available prophylactic for seasonal influenza epidemics^21–24^. To address whether seasonal influenza vaccination can induce cross-reactive antibody responses to bovine A/H5N1 NA, pre and post vaccination levels of NA binding antibodies, NAI antibodies and neutralizing antibodies were determined. It has been shown that although seasonal vaccination minimally increases detectable NA-specific responses, it often induces robust neutralizing antibody (nAb) responses against the strains included in the vaccine formulation, primarily targeting the immunodominant HA head^21,22,24,24–26^. Unsurprisingly, seasonal vaccination did not significantly increase binding IgG responses against either human N1 NA or bovine N1 NA (**Figure 2a-b**). ELLA revealed that vaccination had a statistically significant yet modest increase in NAI antibodies to human N1 NA, but no significant increases in NAI antibodies against bovine N1 NA (**Figure 2c-d**). It should be noted that higher NAI titers post-vaccination may be partly influenced by HA-specific responses^27^. Nonetheless, vaccination did not appear to boost cross-reactive NAI responses, which is consistent with binding IgG responses (**Figure 2a-d**).

**Figure 2.**
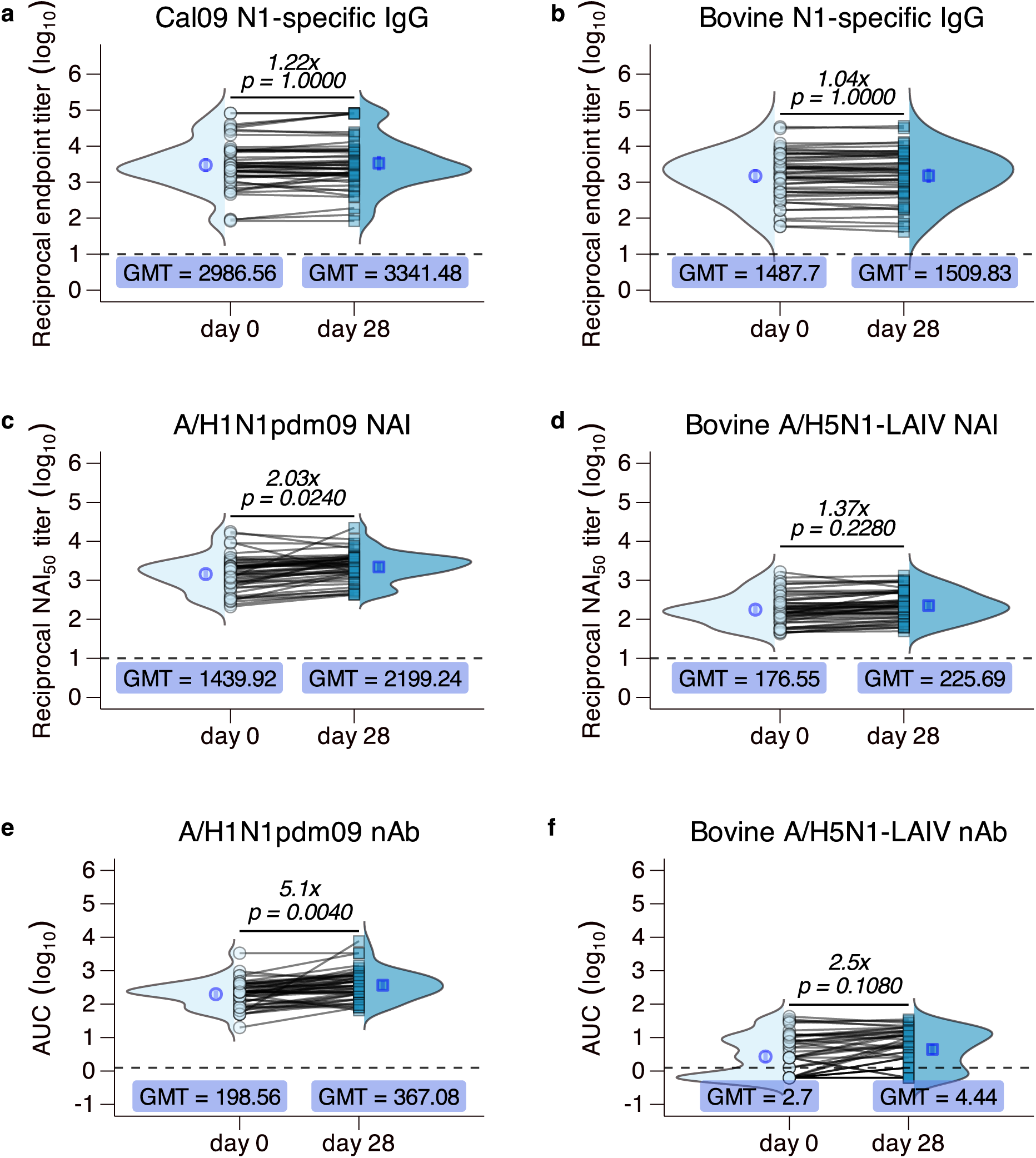
Vaccination does not induce cross-reactive NA-specific responses. (n = 50) Seasonal vaccine recipients were enrolled on the date of seasonal influenza vaccine receipt, and 25 patients were selected at random per each sex, stratified by age group. Serum was taken at the time of enrollment (indicated as “day 0,” indicated by light blue circles) and approximately 28 days later (indicated as “day 28,” indicated as turquoise squares), and subsequently used to evaluate A/H1N1 NA and A/H5N1 NA (**a-b**) total binding IgG via enzyme-linked immunosorbent assay (ELISA) and (**c-d**) NA activity inhibiting (NAI) responses via enzyme-linked lectin assay (ELLA) and (**e-f**) neutralizing antibody responses against an A/H1N1-like virus and bovine A/H5N1-LAIV virus. Dotted lines represent the assay lower limit of detection. Each symbol represents the arithmetic mean of two biological replicates for each sample. Lines between symbols indicate paired baseline and convalescent samples for a single patient. Arithmetic average fold-change values are indicated for each panel, and p-values were generated by paired Wilcoxon signed-rank test. Blue open symbols and error bars represent the geometric mean titer (GMT) and geometric 95% confidence interval (CI), respectively. Ridge plots represent the total distribution of all datapoints.

While seasonal vaccines are formulated specifically with the aim of increasing neutralizing antibody titers against the HA protein, it remained possible that antibody responses boosted by vaccination might lead to an increase in bovine H5 HA neutralizing antibody responses. While nAb against A/H1N1pdm09 increased significantly after vaccination, there was no corresponding increase in A/H5N1-LAIV nAb responses (**Figure 2e-f**). At day 28 post-vaccination, 13 of the initial 24 vaccinees who were seronegative remained seronegative, and 2 vaccinees saw a reduction in nAb response to below the limit of detection (**Figure 2f**). When bovine H5 HA binding antibodies were measured, there was no significant increase detected post vaccination (**Supplemental Figure 3a**). Altogether, the data indicate that current seasonal vaccine formulations do not induce significant levels of binding or functional antibodies that recognize bovine H5 HA or N1 NA proteins.

### Boosting of A/H5N1 cross-reactive antibodies by seasonal IAV infection

Animal studies have suggested infection with human seasonal A/H1N1 viruses can provide partial or complete protection against an A/H5N1 challenge^28–31^. Unlike seasonal vaccination, IAV infection is known to reliably induce NA-specific antibody responses^18,23,27,32,33^. NA-specific antibody responses were measured in our infection cohort (**Figure 3**). For the 16 patients who had sequence-confirmed A/H1N1 infections, Cal09 NA- and bovine A/H5N1 NA-specific IgG were both substantially boosted by infection, albeit to different magnitudes (**Figure 3a-b**), with a 151-fold increase in Cal09 N1 NA binding antibodies and a 55-fold increase in bovine A/H5N1 NA binding antibodies. For A/H3N2-infected patients, there was no consistent trend in binding IgG against both human and bovine N1 NA (**Figure 3a**). These data suggest that A/H1N1 but not A/H3N2 infection boosts bovine N1 NA-specific IgG responses.

**Figure 3.**
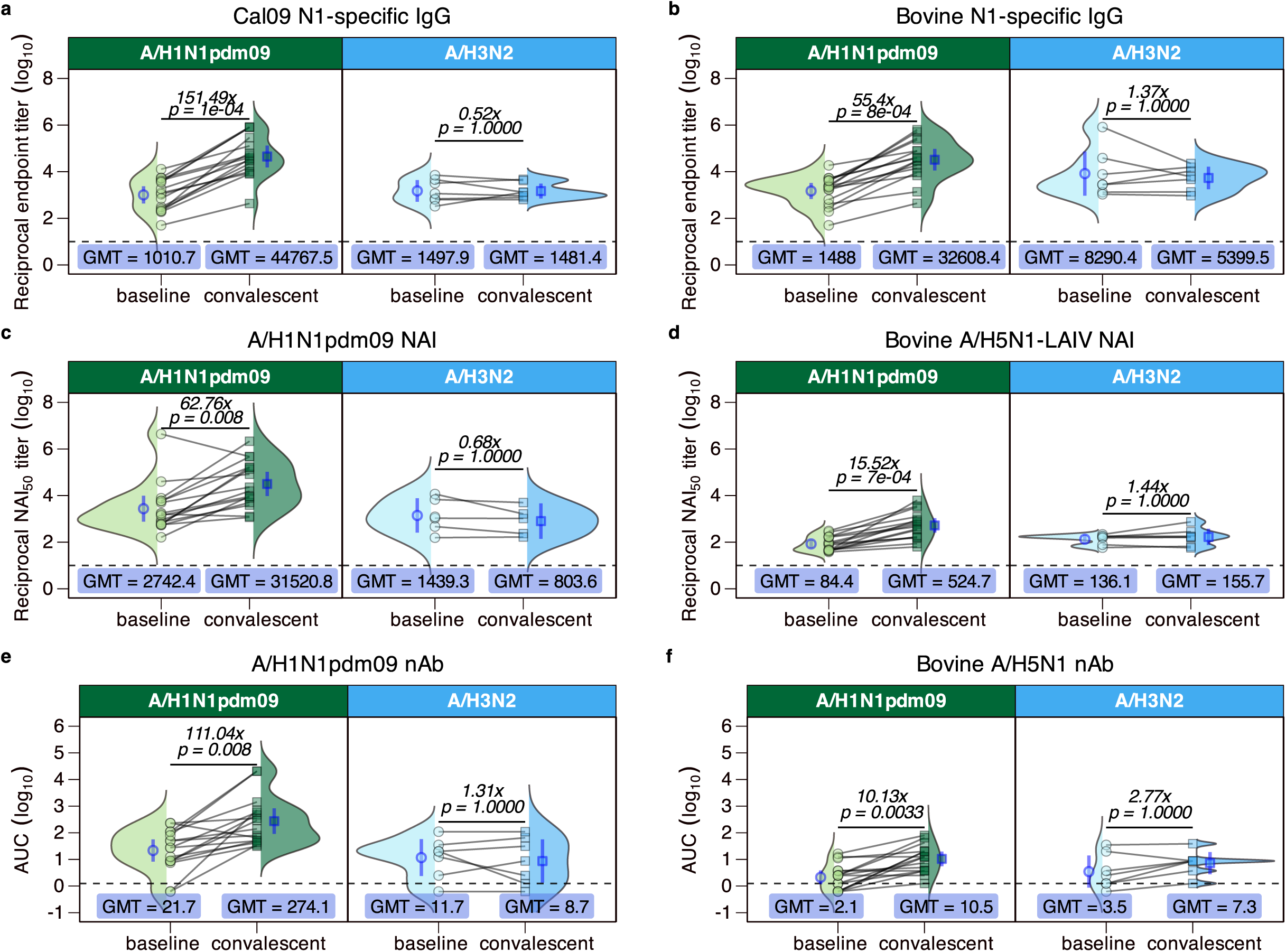
Seasonal A/H1N1 infection induces heterosubtypic NA-specific antibody responses to bovine A/H5N1 NA. (n = 23) Patients presenting to JHH emergency department with ILI and PCR-confirmed Influenza A Virus infection of either A/H1N1subtype (n = 16, green) or A/H3N2 subtype (n = 7, light blue) were used to investigate NA-specific responses. Serum was taken at the time of enrollment (indicated as “baseline,” shown as circles) and approximately 28 days later (indicated as “convalescent,” shown as squares), and subsequently used to evaluate A/H1N1 NA and A/H5N1 NA (**a-b**) total binding IgG via enzyme-linked immunosorbent assay (ELISA) and (**c-d**) NA activity inhibiting (NAI) responses via enzyme-linked lectin assay (ELLA) and (**e-f**) neutralizing antibody responses against an A/H1N1-like virus and bovine A/H5N1-LAIV virus. Grey lines represent the assay lower limit of detection. Dotted lines represent the assay lower limit of detection. Each symbol represents the arithmetic mean of two biological replicates. Lines between symbols indicate paired baseline and convalescent samples for a single patient. Arithmetic average fold-change values are indicated for each panel, and p-values were generated by paired Wilcoxon signed-rank test. Blue symbols and error bars represent the geometric mean titer (GMT) and geometric 95% CI, respectively. Ridge plots representing the total distribution of all datapoints.

The trends we observed in NA binding IgG for A/H1N1-infected patients were recapitulated in NAI responses (**Figure 3c-d**). All but 1 of 16 patients increased in A/H1N1 NAI antibodies and the fold increase was 62x (**Figure 3c**). All 16 patients had a significant increase in A/H5N1-LAIV NAI responses post-infection but with a lower fold-increase of 15.52x (**Figure 3d**). A/H3N2 infected individuals did not show significant increases in NAI responses to human or bovine H1N1 viruses (**Figure 3c-d**). There were also significant increases in nAb titers to both A/H1N1pdm09 and bovine A/H5N1 viruses after A/H1N1 infection but not after A/H3N2 infection (**Figure 3e-f**). In addition to the increase in NAI and nAb responses to bovine A/H5N1-LAIV in the A/H1N1-infected group, there was a significant rise in bovine H5 HA binding IgG titers, where binding IgG titers increased an average of 10.34x by convalescence (**Supplemental Figure 3b**). Altogether, A/H1N1-infected patients were the only group to see an increase in both HA and NA-specific cross-reactive responses against bovine A/H5N1 HA and NA proteins.

### Investigating the contribution of heterotypic N2-specific antibodies to total A/H5N1 NA binding responses

Given the relatively high degree of conservation with respect to structural and functional domains of the NA protein, we determined whether heterotypic antibody to the N2 NA may contribute to the detectable binding IgG at baseline. Pan-NA mAbs often target domains of NA that are essential for either enzymatic function or overall structure of the NA tetramer^34–37^. To address whether N2-targeting IgG are contributing to the observed baseline bovine NA IgG, serum from an age- and sex-stratified subset of our vaccine cohort was depleted of N2-binding IgG (**Supplemental Figure 4).** We used a representative A/H3N2 recombinant NA derived from a locally circulating isolate in Baltimore, MD for all N2-serum depletions. On average, sera showed binding IgG titers that were 3-fold higher than N2-depleted sera against bovine NA (**Supplemental Figure 4**). This reduction occurred in all but one of 25 samples (**Supplemental Figure 4**). These findings suggest that heterotypic N2-specific responses may partially contribute to observed baseline bovine NA antibody responses and substantiate further investigation of cross-group neuraminidase antibody responses.

## DISCUSSION

A key component of risk assessment for A/H5N1 HPAI is a comprehensive understanding of pre-existing immunity at the population level ^38–40^. In the presented work, we illuminate the distinct roles of seasonal human IAV infection and vaccination in shaping baseline cross-reactive antibody repertoire to the neuraminidase (NA) of A/H5N1 viruses. The majority of candidate pandemic HPAI vaccines against A/H5N1 are centered around the HA glycoprotein^17,41–45^. Similarly, available reports detailing the pre-existing antibody repertoire against A/H5N1 are focused on the detection of cross-reactive neutralizing responses which target HA^28,45–48^. Antibody responses against NA are often not neutralizing in function^18,49–53^. While H5 HA is structurally and antigenically distinct from human seasonal H1 HA, they are part of the same phylogenetic group—group 1—and share some identity in sequence, particularly within the more conserved HA stem^48,54–56^. Given the immunodominance of HA during both vaccination and infection, there is a possibility that H5-specific responses at baseline are contributing to the detectable NAI response to bovine A/H5N1-LAIV ^27^, and might also be neutralizing. Other studies have detected low nAb responses against A/H5N1 clades^19,45–47^. Nonetheless, it has long been established that NA-targeting antibodies are an important mediator of protective immunity against IAV, albeit these responses are often not captured in traditional serological assessments like neutralizing antibody assays and hemagglutination inhibition (HAI) assays^18,32,33,51,57–60^. Therefore, N1 NA acting as a shared target between human seasonal A/H1N1 viruses and A/H5N1 HPAI likely allows for the detectable cross-reactive NA-targeting responses we and others report^17,19,45,47^.

Our work describes pre-existing immunity in a population of healthcare workers from Johns Hopkins Hospital (JHH) (**Figure 2**). Understanding the level of baseline immunity against A/H5N1 in this population is essential when considering pandemic potential of A/H5N1, and who is most at risk of occupational or incidental exposures—with healthcare workers being at high risk only second to dairy and poultry farmers who have direct contact with infected animals. Because vaccination is mandated each season for JHH employees, this cohort allows us to investigate the vaccine-induced antibody repertoire as it pertains to cross-reactive bovine A/H5N1 HPAI responses without sacrificing the heterogeneity present at the population level, which is often lost in animal studies^28,29,61,62^. Among healthcare workers, baseline binding IgG against Cal09 NA and against bovine A/H5N1 NA were similar to those among our infection cohort at baseline. When considered with our correlation analyses, binding IgG against both Cal09 NA and bovine A/H5N1 NA appeared to be positively correlated with birth year, although this was not statistically significant for bovine A/H5N1 NA (**Figure 1a**). This trend was consistent for NAI responses, but we saw a slight negative correlation for nAb responses to bovine A/H5N1-LAIV (**Figure 1b-c**). When considering A/H5N1 HA responses, the slight negative correlation with birth year and nAb titer was more exaggerated between birth year and bovine A/H5N1 HA-binding IgG (**Supplemental Figure 2a**). These data highlight important factors and considerations that underlie observed heterogeneity in pre-existing immunity to A/H5N1, such as primary influenza exposures and original antigenic sin^63^.

For seasonal influenza infections, A/H1N1 infection was clearly correlated with the induction of cross-reactive nAb and bovine N1 NA antibodies (**Figure 3**), which was absent in A/H3N2 infected individuals. At least 25% of our infection cohort reported no receipt of a seasonal influenza vaccine in any of the past 5 seasons, and 11 of 23 had not received the seasonal vaccine during the season that they were infected (**Table 2**). This suggests that any differences in baseline NA-specific IgG are unlikely a consequence of vaccine-induced immune responses. However, it is worth noting that the size of our A/H3N2-infected cohort is limited, as the 2023-2024 NH influenza season was dominated by A/H1N1 viruses^64^. Thus, we plan to extend this work to the most recent (e.g., the 2024-2025) NH season and to future influenza seasons, to increase the sizes of both our A/H1N1- and A/H3N2-infected cohorts. Furthermore, while we detect increases in bovine H5 HA (**Supplemental Figure 3c**) and N1 NA (**Figure 3b**) antibody responses after infection, these levels are much lower than those targeting human seasonal HA or NA proteins. Since there are no established correlates of protection from A/H5N1 infection in the human population, we cannot make any conclusions about how these bovine A/H5N1 antibody responses can modulate infection or disease severity. Nonetheless, our data merit additional study for defining immune correlates of protection for both HA- and NA-targeting responses.

Seasonal vaccination did not change H5 binding IgG titers or bovine A/H5N1-LAIV nAb titers, which is expected due to the immunodominance of the globular HA head in vaccine induced antibody responses^21,22,25,65,66^. This corroborates previously published data which indicates that despite no known exposures, the cross-reactive antibodies to H5 HA are detectable, albeit minimal^19,45,47^. While NA has been increasingly considered as a potential immunogen for universal influenza vaccines, our current understanding of how NA-specific responses can mediate immunity are limited. Current practices for seasonal vaccine development and production do not include quantifying NA content^21,65^, despite its role as the common denominator between human influenza viruses and several zoonotic influenza viruses (such as but not limited to: swine A/H1N1, swine A/H1N2, swine A/H3N2, canine A/H3N2, avian A/H1N1, avian A/H2N2, avian A/H3N2, avian A/H5N1, avian A/H9N2, etc.). We show the relevance of A/H1N1 infection, as only A/H1N1 infection was capable of significantly boosting binding, neutralizing, and NA-inhibiting antibody responses against bovine A/H5N1 (**Figure 3**). Relative to HA, NA is a slower moving antigenic target and has lower mutational plasticity^21,60,65^. It must maintain its vital enzymatic function for productive influenza virus infections, which provides a common target across not only IAVs, but also extends to influenza B viruses (IBVs). Despite the discovery of several pan-NA mAbs which have shown protection in lethal challenge models against a panel of human and zoonotic IAVs^18,32,34,35,37,67–69^, NA remains comparatively understudied and overlooked as a viral and vaccine antigen for broadly protective immunity^57,59,60,70^. We show evidence that supports changing current seasonal vaccine formulations to either include greater NA content, or to manipulate immunogen design to increase immunofocusing toward immuno-subdominant domains of IAV HA and NA^25,65,71,72^. The work presented here reiterates the importance of NA as a conserved antigen between human seasonal viruses and A/H5N1 HPAIs and underscores the need for investigation of NA-mediated antibody responses and their role in protective immunity. NA-centered vaccine design would enable robust boosting of cross-reactive N1 antibodies and may serve as a more feasible approach to increasing population-level pre-existing antibodies to A/H5N1 compared to HA-focused vaccines.

## METHODS

### Human subjects enrollment, sampling, and data collection

Serum used in this study was obtained from healthcare workers recruited during the annual Johns Hopkins Hospital employee influenza vaccination campaign in the Fall of 2024 by the Johns Hopkins Centers for Influenza Research and Response (JH-CEIRR). Pre-(immediately prior to vaccination) and post (∼28 day) vaccination human serum was collected from subjects, who provided written informed consent prior to participation. Patients were enrolled at the Johns Hopkins Medical Institute (JHMI) Department of Emergency Medicine or on inpatient floors. Symptomatic patients in the emergency department were screened and tested for influenza from triage by clinical providers using a validated clinical decision guideline tool. Serum was collected at the time of presentation and approximately 28 days later. The Johns Hopkins School of Medicine Institutional Review Board approved this study, with vaccine enrollments under IRB00091667 and acute infection under IRB00306448.

### Viruses

Virus isolates used in this study are as follows; A/Victoria/4897/2022 (A/H1N1 pandemic09-like, 2024-2025 NH vaccine virus, courtesy of John Steel, CDC). Recombinant viruses used in this study include; A/Baltimore/R0675/2019 (A/H1N1 pandemic09-like, GISAID Accession no., EPI_ISL_17617226) and an LAIV-like virus expressing the HA and NA segments of bovine A/H5N1 (GISAID Accession no., EPI_ISL_19014384), which were used for ELLA quantification of NAI antibody and NT_50_ assay for detection of nAb responses. A/Baltimore/R0675/2019 was generated by a 12-plasmid reverse genetics system for the generation of influenza A viruses^73^. Transfection of co-cultured MDCK-SIAT1 and HEK-293T cells with plasmids encoding each of 8 segments belonging to A/Baltimore/R0675/2019, and 4 helper plasmids encoding the polymerase complex of influenza viruses was conducted as previously described^74,75^. Recombinant viruses expressing the HA and NA segments of A/Bovine/Texas/24-029328-01/2024 (GISAID Accession no., EPI_ISL_19014384) were generated as follows; the multibasic cleavage site in HA was mutated to delete the RRKR motif (amino acid positions 342 to 346) to make the virus dependent upon exogenous trypsin^76^. HA and NA segments from A/Bovine/Texas/24-029328-01/2024 were cloned into an 8-plasmid expression system with bi-directional promoters (courtesy of Dr. Seema Lakdawala, Emory University, and Dr. Valerie Le Sage, University of Pittsburgh)^74,75^. The 6 remaining gene segments encoding the internal genes of A/Ann Arbor/6/1960(H2N2) (GISAID Accession no., EPI_ISL_130415), a cold-adapted virus (live-attenuated, indicated as LAIV), cloned into the 12-plasmid reverse genetics system were co-transfected with the bovine-derived A/H5N1 HA and NA described above, with co-cultured MDCKI and HEK-293T cells, as previously described^73–75,77,78^.

### Cell Lines and maintenance

Madin Darby Canine Kidney (MDCK) derivatives, MDCKI and MDCK-SIAT1 (courtesy of Dr. Scott Hensley, University of Pennsylvania) were maintained in cell culture in complete media - hereafter referred to as CM – consisting of Dulbecco’s modified eagle medium (DMEM, Gibco) supplemented with 10% fetal bovine serum (FBS, Gibco), 100 Units/ml Penicillin/Streptomycin (Life Technologies), and 2mM glutaMAX (Gibco). Cells were passaged by washing 2x with PBS (1X, Life Technologies), followed by treatment with trypsin-EDTA (0.5%) (Gibco) and incubation at 37°C for up to 15 minutes, at which point cells had detached. Trypsin was quenched by the addition of an equal volume of CM. Cells were either subsequently passaged or were plated to be used for: virus propagation, virus quantification by tissue culture infectious dose 50% (TCID_50_) assay, and Neutralizing Titer 50% (NT_50_) assay.

### Recombinant neuraminidase HA and NA protein

Recombinant bovine A/H5N1 HA (derived from A/dairy cow/Texas/24-008749-002-v/2024 strain) was obtained from SinoBiologicals (SinoBiologicals, cat no. 41036-V08H). Protein was reconstituted per manufacturer’s instructions to 1 mg/mL and stored at -80°C until use. NA sequences were designed as previously described^79,80^. Briefly, the cytoplasmic, transmembrane, and stalk domains of wild type NA were replaced with an N-terminal signal sequence, 6X-His tag, a tetramerization domain from the human vasodilator-stimulated phosphoprotein (VASP), a thrombin cleavage site, and linker sequence followed by NA sequence^81^. NA constructs were expressed in Expi293F cells and purified by Ni-NTA chromatography as previously described^82^.

### Enzyme-linked immunosorbent assay (ELISA) for quantification of antigen-specific IgG

Antigen-specific IgG was quantified by ELISA as previously described^83,84^. Protein was diluted in 1X PBS (Life Technologies) to 1 µg/mL for all neuraminidase constructs and to 0.5 µg/mL for bovine H5 hemagglutinin (Sino Biological), added to Nunc MaxiSorp 96-well plates (ThermoFisher), and incubated at 4°C for 16 hours. Plates were washed with 1X PBS supplemented with 0.1% Tween-20 (Sigma), designated as PBST. Heat-inactivated serum samples were serially diluted 4-fold, 8 times in blocking buffer, which was comprised of PBST + 5% skim milk (ThermoFisher). Diluted sera was added to plates in duplicate and incubated at room temperature for 1 hour. Goat anti-Human IgG (Gamma chain) Cross-Adsorbed horseradish peroxidase (HRP)-conjugate was used as a secondary antibody. TMB substrate (ThermoFisher) was added to all wells and incubated in the dark for 18-20 minutes. The reaction was stopped by the addition of 0.16 M sulfuric acid, and plates were read at OD_450_ and OD_650_ with background subtraction. Reciprocal endpoint titers were determined as the sera dilution that yielded signal 4x that of secondary antibody alone.

### Enzyme-linked lectin assay (ELLA) for quantification of NAI antibody responses

ELLA was used to quantify neuraminidase activity inhibiting (NAI) antibody responses. In brief, Nunc 96-well Immulon 4 HBX (ThermoFisher) were coated with fetuin from FBS (Sigma) diluted in 1X PBS to 2.5 µg/well. Plates were sealed and stored at 4°C for 12-16 hours. Sera were serially diluted 4-fold, 8 times in assay buffer, which consisted of 0.2 % Tween-20 (Sigma), 1 % BSA (Sigma), 0.1 mg/mL MgCl_2_, and 0.2 mg/mL CaCl_2_, diluted in 1X PBS. A/H1N1 or A/H5N1-LAIV was appropriately diluted in assay buffer and added to serially diluted serum and incubated at 37°C for 1 hour. During the 1 hour incubation, fetuin-coated plates were washed 3x with PBST. Sera/virus mixtures were added to washed plates in duplicate, and each plate contained 8-wells with only assay buffer (to represent no NA activity, or 100% NAI) and an additional 8-wells with virus alone and no antibody (to represent full NA activity, or 0% NAI). Plates were then sealed and incubated for 16 to 18 hours at 37°C/5% CO_2_. Plates were carefully washed with PBST and incubated at room temperature for 2 hours with HRP-conjugated lectin from *Arachis hypogaea*, also referred to as HRP-conjugated peanut agglutinin (PNA, Sigma). Plates were washed a final time and then reacted with SigmaFast OPD substrate (Sigma) away from direct light for approximately 10 minutes. Reactions were stopped by the addition of an equivalent volume of 1N H_2_SO_4_, and OD was read at 490 nm to determine relative NAI.

### Neutralizing Titer 50% (NT_50_) assay for quantification of neutralizing antibody (nAb) responses

NT_50_ assays were conducted as previously reported^85^. Human sera obtained as described above was treated with lyophilized receptor destroying enzyme II (RDE, Hardy Diagnostics) per manufacturer’s instructions. For A/H1N1 and A/H5N1-LAIV viruses, MDCK-SIAT1 cells (courtesy of Dr. Scott Hensley) and MDCKI cells were seeded in complete media (CM) in 96-well plates (Celltreat) 2 days prior to infection. RDE-treated sera were serially diluted in Dulbecco’s modified eagle media (DMEM) supplemented with100 Units/ml Penicillin/Streptomycin (Life Technologies), 2 mM glutaMAX (Gibco), 0.3% bovine serum albumin (Sigma), and 1 or 5 µg/mL of N-acetylated trypsin (Sigma) for assays conducted on MDCK-SIAT1 cells or on MDCKI cells, respectively. 100 TCID50 was added to each well of serially diluted serum and incubated at 33°C/5% CO_2_ for 1 hour prior to infecting cells. Sera and virus mixture was added to cells in quadruplicate and incubated at 33°C 5% CO_2_ for 24 hours. After 24 hours, all plates were washed twice with 1X PBS supplemented with CaCl_2_ and MgCl_2_, and media was replaced before plates were returned to 33°C/5% CO_2_. 120 hours later, plates were fixed with 10% Neutral Buffered Formalin (Leica) and stained with naphthol blue-black for subsequent interpretation. Area under the curve was calculated using GraphPad Prism 10.4.2. Curves were generated by entering the fraction of all four wells per sample protected at each dilution factor. The limit of detection (LOD) was determined to be the smallest possible AUC value generated, e.g., when only one of four wells is protected at the first dilution in the full dilution series. Any samples that had no detectable neutralizing responses were set to be equal to ½ of the assay LOD for use in calculating geometric mean titers (GMTs).

### Serum depletion of antigen-specific antibody

Baseline human sera was depleted of antigen-specific antibody by coupling his-tagged recombinant A/H3N2 NA tetramers from A/Baltimore/R0145/2017(H3N2) (Genbank: MH637451) to magnetic His dynabeads (Invitrogen; ThermoFisher) as previously described^84^. Briefly, 5 µL of heat-inactivated (HI) sera was incubated with 195 µL of N2Baltimore2017 diluted in PBS, for a total of 200 µL, at room temperature with constant agitation for 1 hour. Washed magnetic beads were re-suspended in 50 µL of PBS, and incubated at room temperature with constant agitation for approximately 30 minutes. Tubes containing the above three components were placed on magnetic strip (Invitrogen; ThermoFisher) and incubated until all beads had precipitated out of solution. The supernatant was collected and subjected to ELISA to confirm the successful elimination of all N2Baltimore2017-specific IgG, and to quantify remaining Bovine A/H5N1 NA-binding IgG.

### Data analysis

All statistical analyses shown were calculated in RStudio (version 2025.05.1+513), using either base R or the RStatix packages^86^. Appropriate tests were run as indicated in figure legends. Area under the curve and non-linear regressions were performed either in Graphpad Prism 10.4.2 or in RStudio. All graphs were generated in RStudio.

## DATA AVAILABILITY

All data are available through the Johns Hopkins Data Repository (doi: )

## ACKNOWLEDGEMENTS

The work was supported by NIH/NIAID 75N93021C00045 to the Johns Hopkins Center of Excellence in Influenza Research and Response (JH CEIRR). We thank the members of the Pekosz laboratory for their constructive and supportive feedback. We also thank Dr. Seema Lakdawala and Dr. Valerie Le Sage for providing plasmids encoding bovine A/H5N1 HA(ΔMBS) and NA to generate recombinant A/H5N1-LAIV viruses.

---

**Supplemental Figure 1.**
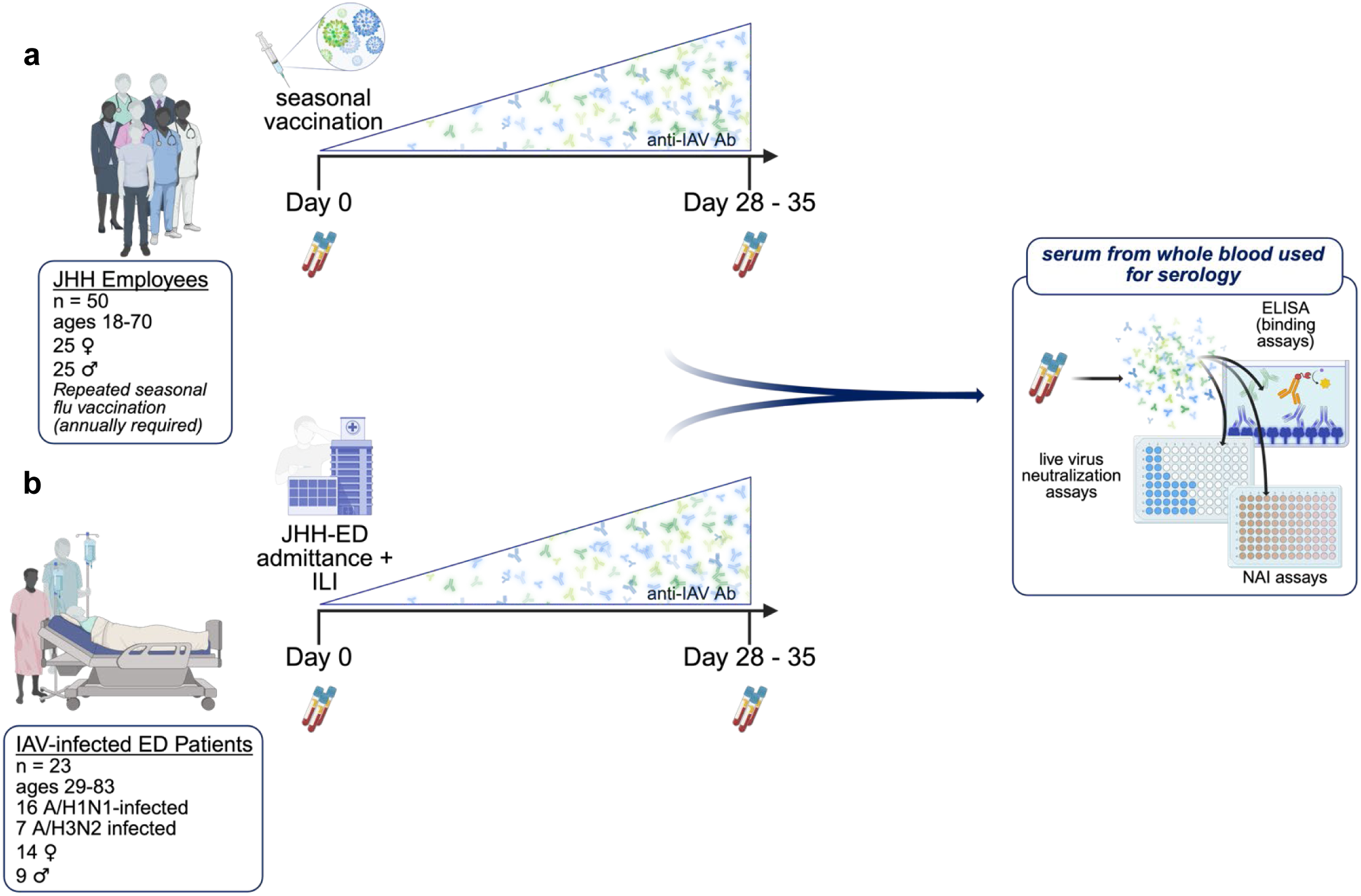
Study design schema. (**a**) vaccine cohort study timeline and composition. (**b**) infection cohort study timeline and composition. Made with Biorender.

**Supplemental Figure 2.**
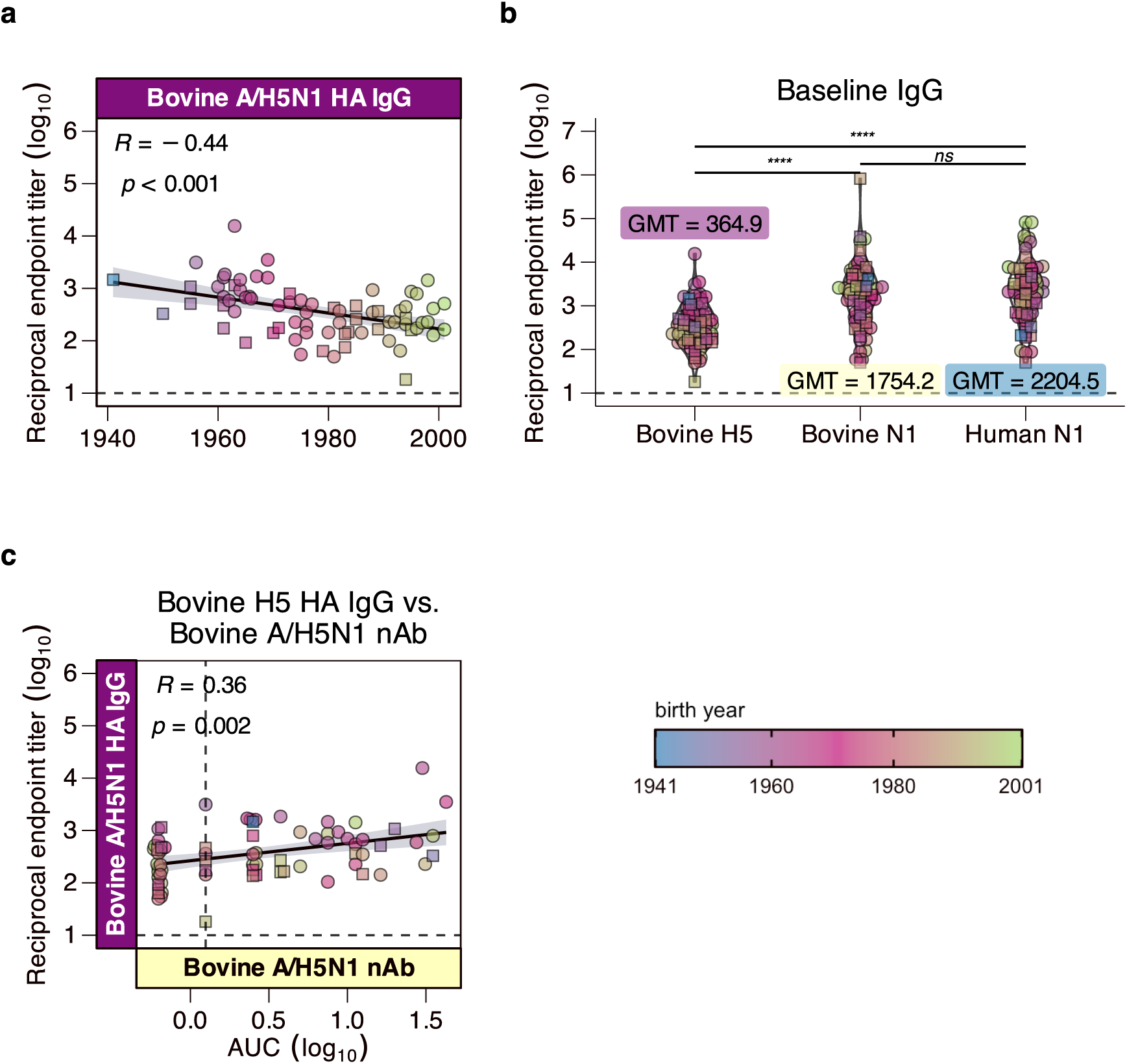
Baseline antibody responses to bovine A/H5N1 hemagglutinin (HA). Baseline binding IgG (**a-b**) against soluble bovine A/H5N1 HA was measured for all samples (n = 73) at the time of enrollment by standard ELISA, plotted by birth year and (**b**) by antigen. Squares represent samples from the infection cohort (n = 23) and circles represent samples from the vaccine cohort (n = 50). (**a**) Spearman coefficient was calculated between baseline binding IgG to A/H5N1 HA and birth year. (**b**) Anti-bovine A/H5N1 HA IgG at baseline was compared to bovine A/H5N1 NA and Cal09 NA binding responses. (**c**) Spearman correlation analyses for binding IgG responses against (**c**) bovine A/H5N1 nAb responses. (**b**) non-parametric one-way ANOVA with Dunn’s post-test shows significantly lower baseline binding IgG titers against bovine A/H5N1 HA than against either NA; *\*\*\*\* = p < 0.0001*, *ns = not significant*. (**b**) geometric mean titer (GMT) is indicated for each group, with H5 binding IgG GMT indicated in purple. (**a, c**) Spearman correlation coefficient and significance are indicated in the upper lefthand corner of each panel. (a-c) Dotted lines indicate the lower limit of detection for the specified assay. Line of best fit is indicated by the black line, with the standard error indicated by the shaded area in each panel.

**Supplemental Figure 3.**
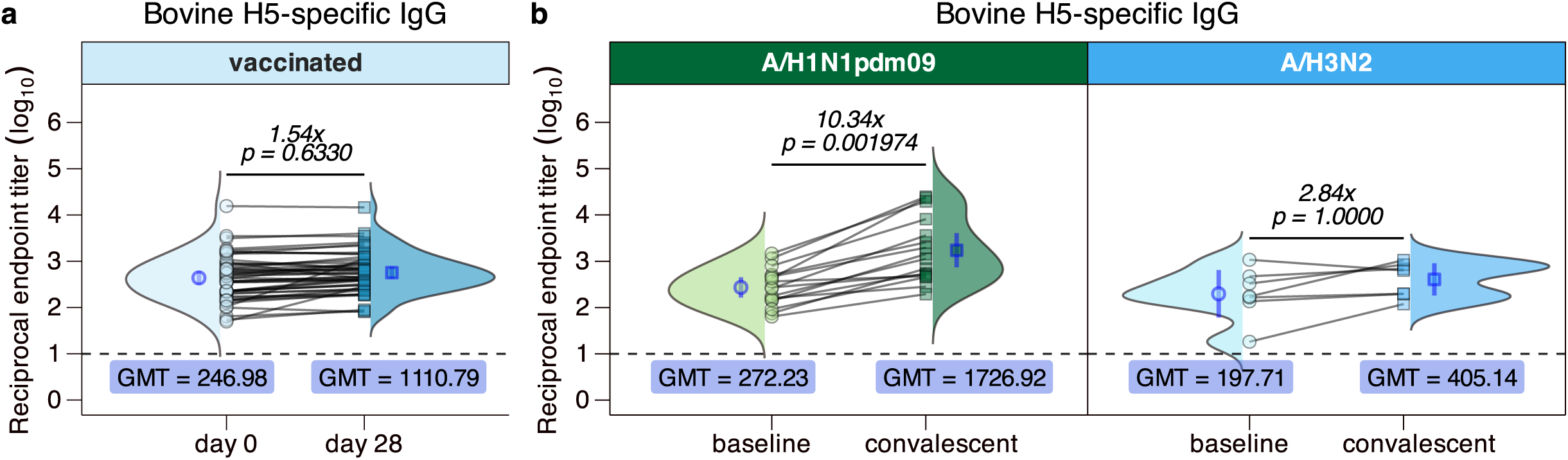
Vaccine- and infection-induced bovine A/H5N1 HA responses. (**a**) serum obtained from vaccine recipients at days 0 (circles) and 28 (squares) post-vaccination were subject to standard binding ELISA to quantify serum IgG against the bovine H5 HA. (**b**) The same quantification of bovine H5 HA-binding IgG was performed on serum from patients infected with circulating (left panel, green) A/H1N1pdm09 or (right panel, blue) A/H3N2 at the time of admittance to the JHH ED (indicated as “baseline,” represented by circles) and approximately 4 weeks later (indicated as “convalescent,” represented by squares). (**a-b**) Dotted lines represent the assay lower limit of detection. Each symbol represents the arithmetic mean of two biological replicates for each sample. Lines between symbols indicate paired baseline and convalescent samples for a single patient. Arithmetic average fold-change values are indicated for each panel, and p-values were generated by paired Wilcoxon signed-rank test. Blue symbols and error bars represent the geometric mean titer (GMT) and geometric 95% CI, respectively. Ridge plots representing the total distribution of all datapoints.

**Supplemental Figure 4.**
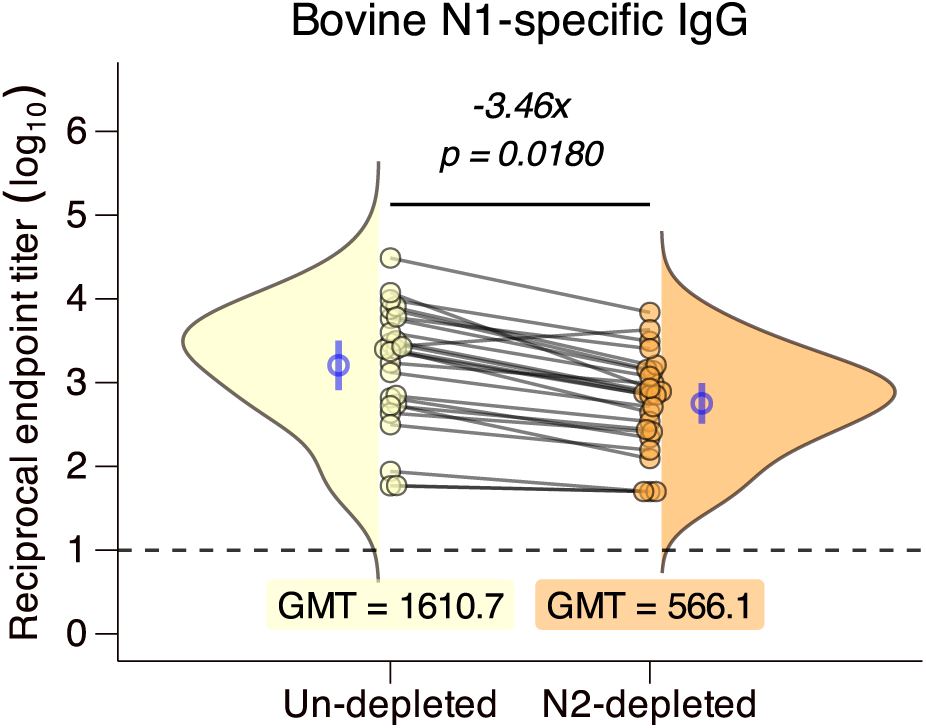
Serum depletion of N2-specific antibody decreases bovine N1 binding IgG. (n = 25) Serum obtained from ½ of the vaccine cohort, stratified by both age and sex, at the time of vaccination was depleted of N2-specific antibody, and subject to ELISA to quantify bovine A/H5N1 NA binding responses. Dotted lines represent the assay lower limit of detection. Each symbol represents the arithmetic mean of two biological replicates for each sample. Lines between symbols indicate paired baseline and convalescent samples for a single patient. Arithmetic average fold-change values are indicated for each panel, and p-values were generated by paired Wilcoxon signed-rank test. Blue symbols and error bars represent the geometric mean titer (GMT) and geometric 95% CI, respectively. Ridge plots representing the total distribution of all datapoints.

